# 4-phenylbutyrate restored GABA uptake and reduced seizures in *SLC6A1* variants-mediated disorders

**DOI:** 10.1101/2021.12.08.471799

**Authors:** Gerald Nwosu, Felicia Mermer, Carson Flamm, Sarah Poliquin, Wangzhen Shen, Kathryn Rigsby, Jing-Qiong Kang

## Abstract

We have previously studied the molecular mechanisms of solute carrier family 6 member 1 (*SLC6A1*) associated with a continuum of neurodevelopmental disorders, including various epilepsy syndromes, autism, and intellectual disability. Based on functional assays of variants in a large cohort with heterogenous clinical phenotypes, we conclude that partial or complete loss of GABA uptake function in the mutant GAT-1 is the primary etiology as identified in GABAA receptor mutation-mediated epilepsy and in cystic fibrosis. Importantly, we identified that there are common patterns of the mutant protein trafficking from biogenesis, oligomerization, glycosylation, and translocation to the cell membrane across variants with the conservation of this process across cell types. Conversely any approach to facilitate membrane trafficking would increase presence of the functional protein in the targeted destination in all involved cells. PBA is an FDA-approved drug for pediatric use and is orally bioavailable so it can be quickly translated to patient use. It has been demonstrated that PBA can correct protein misfolding, reduce ER stress, and attenuate unfolded protein response in neurodegenerative diseases, it has also showed promise in treatment of cystic fibrosis. The common cellular mechanisms shared by the mutant GAT-1 and the mutant cystic fibrosis transmembrane conductance regulator led us to test if PBA and other pharmaco-chaperones could be a potential treatment option for *SLC6A1* mutations. We examined the impact of PBA and other small molecules in a library of variants and in cell and knockin mouse models. Because of the critical role of astrocytic GAT-1 deficit in seizures, we focused on astrocytes, and demonstrated that the existence of the mutant GAT-1 retained the wildtype GAT-1, suggesting aberrant protein oligomerization and trafficking caused by the mutant GAT-1. PBA increased GABA uptake in both mouse and human astrocytes bearing the mutations. Importantly, PBA increased GAT-1 expression and suppressed spike wave discharges (SWDS) in the heterozygous knockin mice. Although the detailed mechanisms of action for PBA are ambiguous, it is likely that PBA can facilitate the forward trafficking of the wildtype GAT-1 favoring over the mutant GAT-1, thus increasing GABA uptake. Since all patients with *SLC6A1* mutations are heterozygous and carry one wildtype functional allele, this suggests a great opportunity for treatment development by leveraging the endogenous protein trafficking pathway to promote forward trafficking of the wildtype in combination with enhancing the disposal of the mutant allele as treatment mode. The study opens a novel avenue of treatment development for genetic epilepsy via drug repurposing.

## Introduction

Genetic variation and the subsequent protein misfolding are a major cause of disease throughout the life-span^1–4^. In childhood, pathogenic variants or mutations in numerous genes give rise to a wide range of neurodevelopmental disorders. GABA transporter 1 (GAT-1) encoding *SLC6A1* belongs to one such gene^5–7^. *SLC6A1* is a member of neurotransmitter subgroups, which include 3 γ-aminobutyric acid (GABA) transporters (GAT), 2 glycine transporters (GLY), and the monoamine (dopamine (DAT), serotonin (SERT) and norepinephrine (NET)) transporters in the solute carrier (SLC6) family^8^. The SLC6 family transporters are integral membrane solute carrier proteins characterized by the Na^+^-dependent translocation of small amino acid or amino acid-like substrates. As aforementioned, mutations in the family are frequently associated with epilepsy, autism, and various neurodevelopmental disorders^5,6,9^. Based on currently reported data, all patients are heterozygous, carrying the mutant allele and one functional wildtype allele, arguing for haploinsufficiency with or without dominant negative suppression from the mutant allele as described in *GABRG2* mutations^10^.

We have previously characterized the functional consequences of both missense and nonsense *SLC6A1* mutations associated with a wide spectrum of disease phenotypes. We identified the common mechanisms of the molecular pathophysiology underlying the heterogenous clinical phenotype. The molecular pathophysiological mechanisms include reduction or loss of GABA uptake, endoplasmic reticulum (ER) retention of the mutant GAT-1 protein, and reduced membrane and total GAT-1 protein expression due to impaired protein trafficking^9,11–13^. The mechanisms directly contributing to diminished GAT-1 function include decreased membrane protein trafficking due to protein misfolding and altered protein stability. This is a common phenomenon also observed in GABA_A_ receptor subunit mutations in our previous studies^14–16^. Protein misfolding has been widely studied for later onset neurodegenerative diseases but much less in the early onset childhood neurodevelopmental disorders such as epilepsy or autism. Our studies in both GABA_A_ receptors and more recently in GAT-1 indicate that ER retention of the mutant protein due to protein misfolding is common among pediatric neurological disorders. Thus leveraging the ER pathway and promoting membrane protein trafficking could be a novel treatment target for genetic epilepsy as well as other related disorders^17^. This could be disease modifying as leveraging the ER pathway via therapeutic intervention could correct the mutant protein upstream of signaling cascades of the complex disease mechanisms.

4-phenylbutyrate (PBA) or 4-phenylbutyric acid is a salt of an aromatic fatty acid. It is used to treat urea cycle disorders, because its metabolites offer an alternative pathway to the urea cycle allowing excretion of excess nitrogen^18^. PBA is a chaperone seen to reduce ER stress^19,20^, while also functioning as a histone deacetylase inhibitor ^21,22^. We have previously identified a trafficking deficient γ2 subunit protein which can modify the epileptic phenotype in mice suggesting its role in the etiology of disease^1,10,23^. Similarly, we recently identified that impaired trafficking and endoplasmic reticulum (ER) retention as the major molecular pathophysiology of mutant GAT-1 proteins. This thus implicates the potential of the same mechanism-based rescue of *SLC6A1* mutants by PBA. PBA has been used to treat cystic fibrosis mutant ΔF508-CF Transmembrane Conductance Regulator (ΔF508-CFTR), by targeting the same pathophysiological mechanism, with clinical trials showing promise^24,25^. The mutation ΔF508-CFTR causes the mutant protein to be unstable and misfold, thus remaining trapped in the ER and unable to reach the cell membrane. PBA can stabilize the mutant CFTR in the ER, allowing it to reach the cell surface, thus rescuing the disease phenotype. Thus, the same PBA treatment paradigm can be utilized in the identified GAT-1 trafficking deficits.

In this study, we evaluated the impact of PBA on eight *SLC6A1* mutations including in patient iPSC derived astrocytes and knockin mice. We tested the rescue effect of PBA in eight representative mutations located in the different functional domains of GAT-1 protein peptide due to either missense or nonsense mutations. We evaluated the impact of PBA on the mature and ER-bound protein expression as well as the GABA reuptake activity. Because of the unique role of GAT-1 in thalamic astrocytes and the associated seizures phenotypes^26–29^, we characterized the impact of PBA on GABA uptake activity in astrocytes derived from patient cells and in knockin mouse models and determined the effect of PBA in two *SLC6A1* mutation knockin mouse lines including using the video monitored synchronized EEG recordings.

## Materials and methods

### *SLC6A1* variants information

The patient variants are selected from the lab cDNA library built based on our previous studies^6,11,30,31^. Those variants are representative for missense and nonsense, variation location in N terminus vs C-terminus, as well as common patterns of function and trafficking defects of the variants.

### *SLC6A1* mutation knockin mouse models

Both mutations *SLC6A1 (A288V)* and *SLC6A1 (S295L)*, for which the knockin mouse models have been created, have been characterized in our previous study^11^. The *SLC6A1 (A288V)* mouse line was generated in collaboration with the university of Connecticut health center UConn Health and the *SLC6A1 (S295L)* mouse line was generated by Shanghai Model Organisms (Shanghai Model Organisms Center, Inc. Cat. NO. NM-KI-190014). Both mouse lines are developed with the CRISPR-CAS9 global knockin approach. Both mouse lines are maintained in the C57BL/6J mice (Jax Stock #000664) and the mice used for biochemistry experiments were between 2-8 months old. The mice used for EEG recordings were between 2-4 months old. The mice used in the study were generated by breeding the heterozygous with the wildtype. Both male and female heterozygous mice were used for breeding of both mouse lines and for experiments.

### Cloning of GABA transporter 1

The plasmid cDNA encoding enhanced yellow fluorescent protein (YFP)-tagged rat GAT-1 has been previously described^32^. The coding region of rGAT-1 was inserted into pEYFP-C1 (Clontech, Palo Alto, CA). QuikChange Site-directed Mutagenesis kit was utilized to introduce the GAT-1 variants into a wildtype GAT-1 plasmid. The product was then amplified via polymerase chain reaction, transformed using DH5α competent cells, and plated. A clone was chosen and grown overnight. All the GAT-1 variants were confirmed by DNA sequencing. Both the wildtype and the variant cDNAs were prepared with Qiagen Maxiprep kit.

### Mouse cortical astrocyte cultures and transfections

Mouse cortical astrocyte cultures and transfection were prepared as previously described. Briefly, the cortices of postnatal day 0-3 pups, were dissected. The tissues were minced after removing the meninges and then digested with 0.25% trypsin for 10 minutes at 37°C. The cells were cultured in poly-L-lysine coated 100 mm^2^ dishes and maintained in Dulbecco’s Modified Eagle’s Medium supplemented with 10% FBS and 1% penicillin/streptomycin. The astrocytes used for experiments were at passage 2. The details for astrocytes culture and transfection protocol are described in supplementary Methods.

### Human patient derived astrocytes

The patient-derived and CRISPR-corrected iPSCs were obtained in collaboration with Dr. Jason Aoto’s lab (University of Colorado). The normal human iPSC clone was purchased from Thermo Fisher (A18945). The detailed iPSC culture, solutions, and differentiation protocols are described in Supplementary Methods. Briefly, the corrected and patient cell lines were maintained in plates coated with Geltrex (1:50 DMEM/F12) overnight using mTeSR (Stem Cell technologies). The media was refreshed daily for iPSCs. The differentiation of neural progenitor cells (NPCs) was induced by STEMdiff SMADi Neural induction kit from STEM Cell Technologies. The differentiation of astrocytes was initiated using the Astrocyte medium (ScienCell) for 27 to 30 days and the cells were passaged at ~70% confluence^33^. The differentiation of astrocytes was initiated at NPC day 5 from passage 1. The experiments in astrocytes were carried out between 27 to 30 days after differentiation from NPCs.

### Radioactive ^3^H-labeled GABA uptake assay

The radioactive ^3^H-labeled GABA uptake assay in HEK293T cells, mouse cortical astrocytes, and human patient derived iPSCs derived astrocytes was modified from the protocol used in our previous studies as detailed in Supplementary Methods. The protocols for GABA reuptake assay in cultured mouse astrocytes or iPSC-derived cells were modified from the GABA uptake protocol on HEK293T cells^11^. Control dishes treated with the GAT-1 inhibitors Cl-966 and NNC-711 and in some cases GAT-3 inhibitor SNAP5114 were included to make sure the radioactive counts measured are specific.

### Measurement of surface and total expression of GAT-1 using flow cytometry

The protocol used for measurement of surface and total expression of GAT-1 using flow cytometry has been described previously for studies on GABA_A_ receptor subunit variants and *SLC6A1* variants ^11,13^. Both the surface and the total GAT-1 protein were probed with rabbit polyclonal anti-Rabbit GAT-1 and 10,000 single cells were evaluated. The empty vector pcDNA mock transfected cells and untransfected cells were included as reference each time.

### Confocal microscopy and image acquisition

Live cell confocal microscopy was performed using an inverted Zeiss laser scanning microscope (Model 510) with a 63 × 1.4 NA oil immersion lens, 2-2.5 × zoom, and multi-track excitation^8^. Cells were plated on poly-I-ornithine and Laminin-coated coverslips or glass-bottom imaging dishes at the density of 1-2 × 10^5^ cells per dish and co-transfected with 0.5 μg of the wildtype or the mutant GAT-1 plasmids with 0.5 μg of the ER marker ER^CFP^ per 35-mm glass-bottomed culture dish with PEI, as per our standard lab protocol. All images were obtained from live cells with single confocal sections averaged from 8 times to reduce noise.

### PBA administration and EEG recordings

Mice of both sexes at 2-4 months old were dosed with PBA (100 mg/kg, i.p.) or vehicle for 7 days. The dose was chosen based on a dose–response experiment performed using increasing doses of PBA from 100 to 800□mg/kg to determine the most efficacious dose^34^. PBA solution was prepared by dissolving PBA in 0.9% normal saline and then titrating equimolecular amounts of 4-phenylbutyric acid (Sigma, Madrid, Spain) and sodium hydroxide to pH 7.4. The working solution was stored at 4 °C.

### Synchronized video-monitoring EEG recordings

Surgery to implant the EEG head mount and the video-monitoring synchronized EEG recordings were conducted as previously described ^8,9^. Synchronized video EEGs were recorded from 8 week to 4 month old C57BL/6J mice one week after electrode implantation with synchronized EEG monitoring system from Pinnacle Technology based on previous study^10^. Briefly, mice were anesthetized with 1–3% isoflurane and four epidural electrodes (stainless steel screws affixed to one head mount) were placed on the brain surface above the bregma and secured in place with dental cement and surgical stiches. EMG leads were inserted into the trapezius muscle. Mice were allowed to recover from the EEG head mount implantation surgery for 5-7 days before EEG recording. The analgesic, ketofen (5mg/kg), was administered 5 min before surgery and for three-days post-surgery. Video-EEG monitoring lasted for 48 hrs. and mice were freely moving during EEG recordings. The Racine Scale was used to identify mouse behaviors such as behavioral arrest, myoclonic jerks, or generalized tonic clonic seizures during the EEG recordings to determine if mice exhibit behavioral seizures. At least 48 hours of baseline EEG recordings were obtained and analyzed for each mouse.

### EEG analysis with Seizure Pro software

For recordings the sampling rate is set at 400 Hz with a pre-amplifier gain of 100 Hz. EEG and EMG channels have a filter set at 25.00. Recording takes place over a 48-hour period and mice are checked daily for health concerns, and food/water access. EEG recordings are scored blind by a skilled scorer using the Sirenia Seizure Pro software. A power analysis is performed using the theta frequency band of 5-7 Hz. An average power is calculated using baseline recordings and applied to seizure analysis for treatment recordings (~1500 uV^2^). Seizures identified by the software were confirmed using video recordings of the period. The Racine scale was used for seizure identification (Stage 1: mouth/facial movements; Stage 2: head nodding; Stage 3: forelimb clonus; Stage 4: rearing; Stage 5: rearing and falling). The presence of spike and wave discharges were identified and compared across treated and non-treated recordings.

### Statistical analysis

Data were expressed as mean ± SEM. Proteins were quantified by Odyssey software and data were normalized to loading controls and then to wildtype subunit proteins, which was arbitrarily taken as 1 in each experiment. Fluorescence intensities from confocal microscopy experiments were determined using MetaMorph imaging software, and the measurements were carried out in ImageJ as modified from previous description ^4, 11^;^8, 9^. For statistical significance, we used oneway analysis of variance (ANOVA) with Newman-Keuls test or Student’s unpaired *t*-test. In some cases, one sample *t*-test, unpaired *t*-test or paired *t test* was performed (GraphPad Prism, La Jolla, CA), and statistical significance was taken as p < 0.05.

### Data Availability

The data supporting the findings of this study are available within the article and its Supplementary Material.

## Results

### *Both missense and nonsense SLC6A1* variants caused partial or complete loss of GABA uptake function due to reduced GAT-1 surface protein expression

*SLC6A1* variants are distributed in various locations of the GAT-1 protein peptide and are associated with various epilepsy syndromes and neurodevelopmental disorders ^6,9,12,13,30,31^. We selected 8 mutations in the study as they are representative of missense and nonsense mutations with premature stop codon mutations generated at the N or C-terminus (Figure 1A). HEK293T cells were transfected with the empty vector pcDNA, wildtype, or the mutant GAT-1^YFP^ for 48 hrs before GABA uptake assay. All mutations had reduced GABA uptake activity (Figure 1B), levels similar to what is observed in cells expressing the wildtype GAT-1 treated with GAT-1 inhibitors Cl-966 (50 μM) or NNC-711 (35 μM) (Figure 2B). Consistently, the cell surface expression of the GAT-1 was reduced in the mutant (Figure 1C). However, the expression of total GAT-1 was reduced in some mutations but not in the D410E and L460R mutations. It is interesting that the mutations resulted in loss of GABA uptake function regardless of missense or nonsense variants in N-terminus or C-terminus (Figure 1D). In all surveyed variants, the GABA uptake was less than 50% of the wildtype. All eight variants had reduced surface expression with varying magnitude as evaluated by a high-throughput assay, flow cytometry (0.002 for E16X, 0.402 for V125M, 0.394 for A288V, 0.248 for S295L, 0.532 for G362R, 0.73 for D410E, 0.75 for L460R and 0.21 for W495X vs wt=1). Most mutations had reduced total protein expression except D410E and L460R (0.01 for E16X, 0.488 for V125M, 0.562 for A288V, 0.536 for S295L, 0.558 for G362R, 0.93 for D410E, 0.98 for L460R and 0.146 for W495X). This suggests there is a correlation of the reduction of surface protein and total protein but no direct correlation of the GABA uptake function with protein as some mutant protein is trafficking competent.

**Figure 1:**
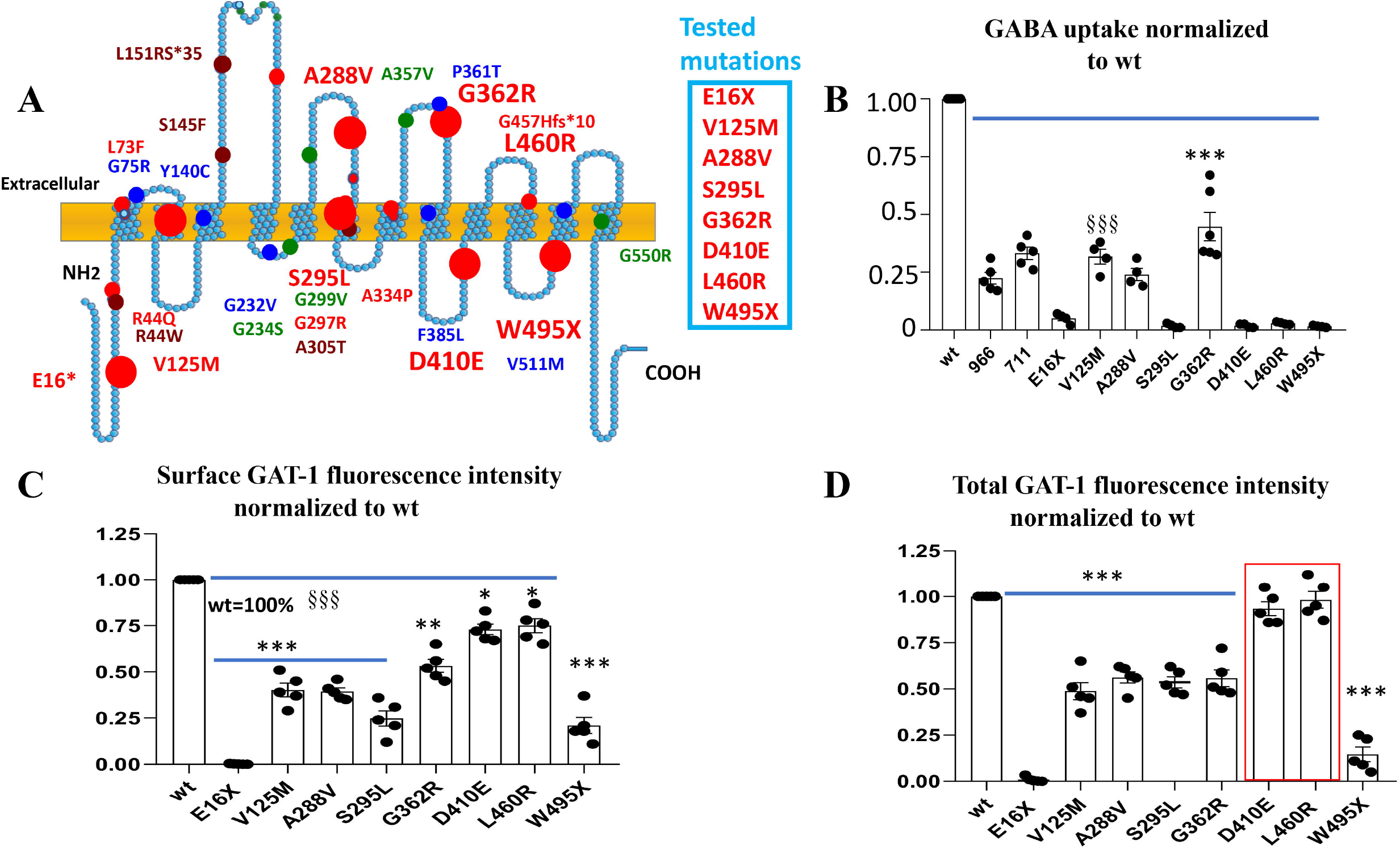
Reduced function and trafficking of the mutant GABA transporter 1 encoded by *SLC6A1* variants associated with epilepsy, autism, ADHD, and intellectual delay. **A**. Schematic presentation of mutant GABA transporter 1 (GAT-1) protein topology and locations of representative mutations in human *SLC6A1* associated with various epilepsy syndromes and neurodevelopmental disorders as described in our previous work. These mutations are distributed in various locations and domains of the encoded GAT-1 protein peptide. The large red dots represent the eight mutations evaluated in the study. (**B, C, D**). HEK293T cells were transfected with the wildtype or the mutant GAT-1^YFP^ for 48 hrs. **B**. The graphs represent the altered GABA reuptake function of the mutant GAT-1 encoded by 8 different *SLC6A1* variants in HEK293T cells measured by the high-throughput ^3^H radio-labeling GABA uptake on a liquid scintillator with QuantaSmart. 966 stands for the wildtype treated with GAT-1 inhibitor Cl-966 (50 μM) and NNC-711 for the wildtype treated with NNC-711 (35 μM) for 30 min before preincubation. **C. D** The flow cytometry histograms depict the relative surface (**C**) or total (**D**) expression of the wildtype and the mutant GAT-1. The relative total expression level of GAT-1 in each mutant transporter was normalized to that obtained from cells with transfection of the wildtypes. (N=4-5 different transfections, δδδ P<0.001 overall mutations vs wt, *P<0.05, **P<0.01, \*\*\**p*< 0.001 vs wt, one-way analysis of variance (ANOVA) and Newman-Keuls test. Values were expressed as mean ± S.E.M.)

**Figure 2.**
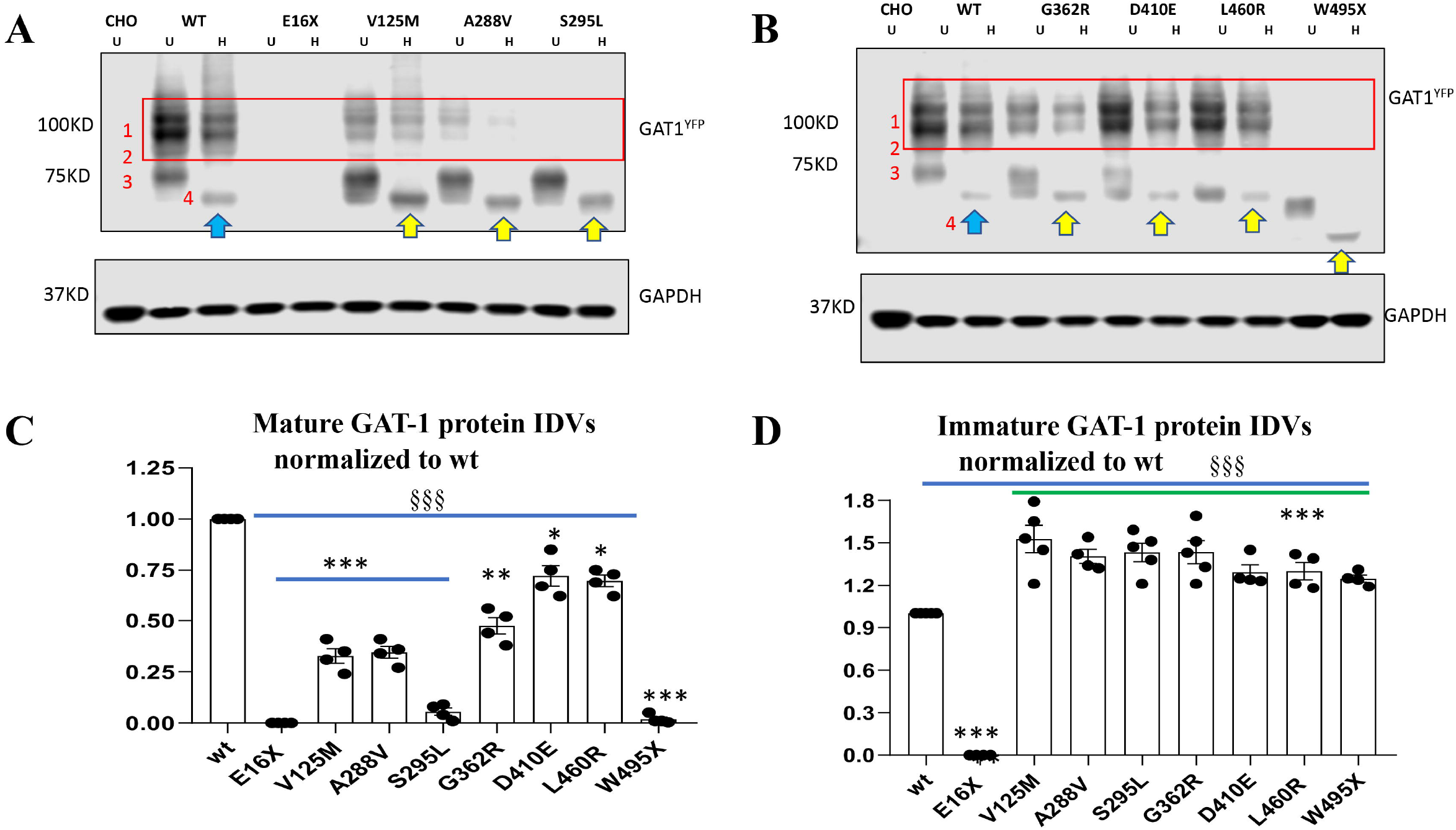
All surveyed mutant GAT-1 transporters had less mature but more immature form of the GAT-1 protein. **A, B.** The total lysates of HEK293T cells expressing the wildtype or variant GAT-1 were undigested (U) or digested with Endo-H (H) and then analyzed by SDS-PAGE. The membrane was immunoblotted with a rabbit anti-GAT-1. The red-boxed region represents the mature form of GAT-1 in cells. (**C**). The graph represents the normalized integrated protein density values (IDVs) of the mature form of GAT-1 defined by being Endo-H resistant normalized to wt mature form of GAT-1 (bands 1+2). (**D**). The graph represents the normalized integrated protein density values (IDVs) of the immature form of GAT-1 defined by being Endo-H unresistant (shifted to a lower level after H digestion) normalized to wt immature form of GAT-1 (band 3 shifted to band 4). (N=4-5 different transfections, δδδ P<0.001 overall mutations vs wt, *P<0.05, **P<0.01, \*\*\**p*< 0.001 vs wt, one-way analysis of variance (ANOVA) and Newman-Keuls test. Values were expressed as mean ± S.E.M.)

### *SLC6A1* mutations caused less mature form but more ER retained immature form of GAT-1 protein

The reduced GABA uptake activity could be caused by reduced cell surface expression or the altered gating kinetics of the transporter channels. Based on our studies on *SLC6A1* and GABA_A_ receptor epilepsy mutations, the reduced cell surface expression of the mutant protein is the major mechanism and is mainly due to ER retention of the misfolded mutant protein, while the altered gating is a minor mechanism. Based on our studies on GABA_A_ receptors, only those proteins that are mature for glycosylation and that have trafficked beyond the ER to reach the cell surface and synapse can function ^4, 12^. We thus determined the maturity of the mutant GAT-1 with Endo-H digestion, which removes the ER added glycan but not the glycan added beyond ER. As reported in our previous study ^2^, the GAT-1^YFP^ protein runs with 3 bands (band 1,2 and 3), 108 KDa, 96 KDa and 90 KDa respectively (Figure 2 A and B). Endo-H treatment removes the glycan added in ER but not those added beyond ER. Thus, the ER retained protein run at a lower molecular mass (band 4). The band 1 and band 2 are classified as mature form of GAT-1 while the band 3 and the down-shifted band 4 are classified as immature GAT-1. The GAT-1(E16X) mutant protein was undetectable likely because of fast degradation of the small peptide due to the early premature stop codon. The GAT-1(W495X) mutation ran at a more reduced molecular mass due to the premature stop codon generated by the nonsense mutation (Figure 2B). Most of the mutant GAT-1 had reduced mature form of GAT-1 (0.00 for E16X, 0.327 for V125M, 0.345 for A288V, 0.05 for S295L, 0.475 for G362R, 0.72 for D410E, 0.69 for L460R and 0.019 for W495X) but increased Endo-H sensitive immature form of GAT-1 (0.00 for E16X, 1.526 for V125M, 1.405 for A288V, 1.432 for S295L, 1.434 for G362R, 1.293 for D410E, 1.30 for L460R and 1.248 for W495X). (Figure 2 C and D), suggesting ER retention of the mutant transporter as in our previous studies on both GABA_A_ receptor and transporters. The mature form of GAT-1 in GAT-1 (V125M, A288V, S295L and S295L and G362R) are correlated with the level of GABA uptake function. The mature form of GAT-1(D410E) and GAT-1(L460R) in HEK 293T cells does not correlate with the GABA uptake function. The GAT-1(E16X) and GAT-1(W495X) had almost no mature GAT-1 protein (Figure 2C). By contrast, the immature GAT-1 as shifted with Endo-H was higher in all surveyed mutant GAT-1 except GAT-1(E16X) as the short, truncated protein peptide in the mutant protein may be subjected to fast disposal inside ER.

### 4-phenylbutyrate (PBA) increased GABA uptake in the wildtype and the mutant transporters in HEK293T cells

We have previously demonstrated that ER retention of the mutant protein can exacerbate the disease phenotype by preventing the efficient trafficking of the wildtype subunits ^9, 13^. We then determined if PBA, a chaperone inducer, can rescue the wildtype and the mutant GAT-1 activity. We first determined the effect of PBA on the wildtype GAT-1. The HEK 293T cells were transfected with GAT-1^YFP^ for 48hrs and PBA was applied with a series of concentrations from 0 to 4 mM ((Figure 3A) and time durations from 0 to 48 hrs (Figure 3B) before evaluation. PBA concentration and time dependently increased the GABA uptake activity (Figure A and B). However, the effect of PBA at 2mM and 24 hrs was the most optimal (1.24±0.05 for 24 hrs and 1.24±0.09 for 4 mM vs wildtype 0) as occasional cell loss was observed in cells treated with PBA at 4 mM or 48hrs. We thus chose PBA 2mM and 24 hrs for the following experiments.

**Figure 3.**
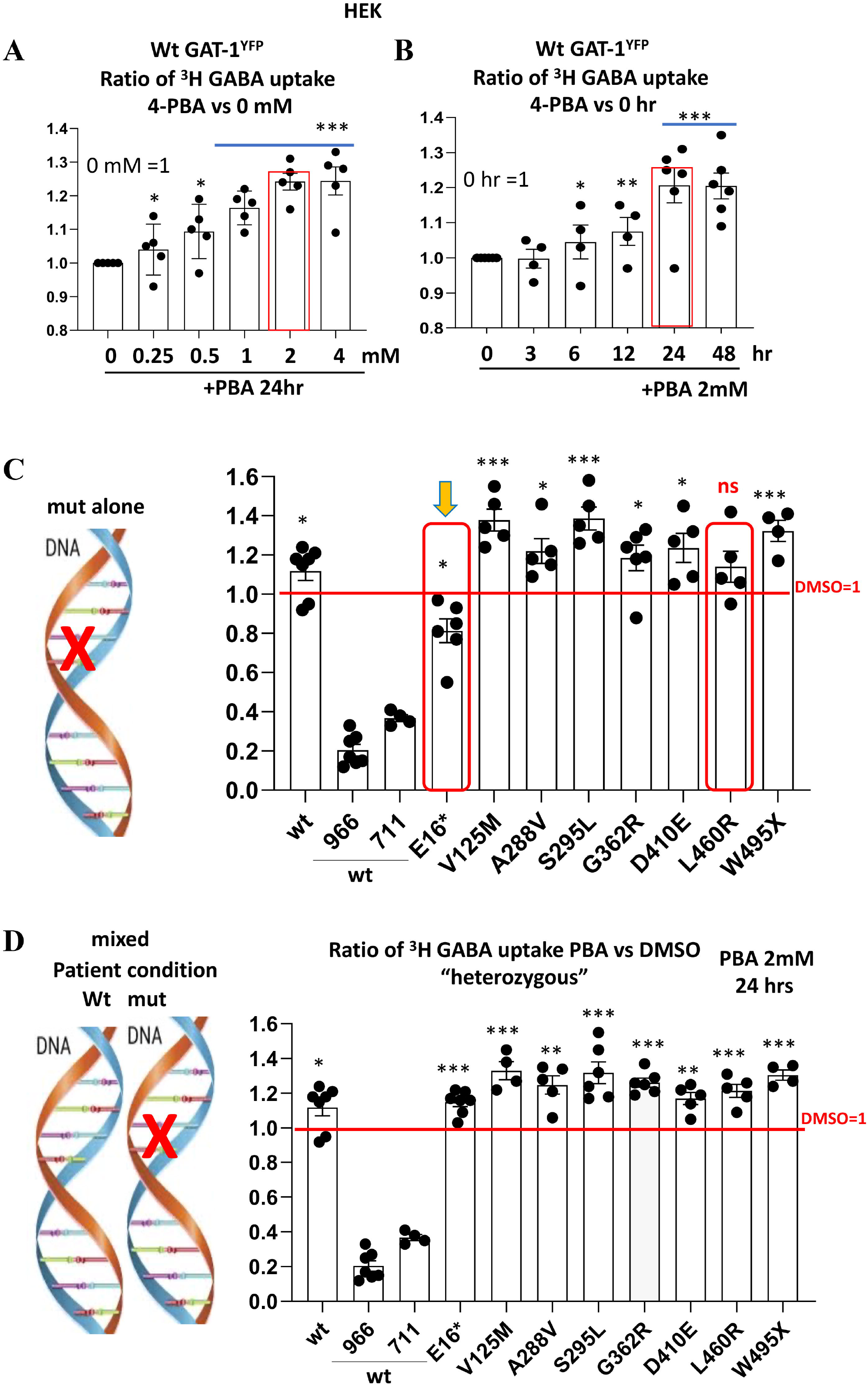
4-Phenylbutyric acid (PBA) dose and time-dependently increased the GABA uptake in cells expressing the wildtype GAT-1^YFP^ in the wildtype alone or in the mutant conditions. **A, B.** HEK293T cells were transfected with wildtype GAT-1^YFP^ (wt) for 48 hr, PBA (2mM) was applied dropwise to each dish at different concentrations (**A**) and incubated for time durations (**B**). PBA from stocking solution (2M) was diluted with DMEM 100 μl to desired concentration. **A**. The graphs represent the altered GABA reuptake function of the wildtype GAT-1 in HEK293T cells treated with PBA for a series of different concentrations. The GABA uptake activity of cells treated with PBA of different concentrations was normalized to the sister cultures treated with DMSO alone for 24 hrs. (**A**). The graphs represent the altered GABA reuptake function of the wildtype GAT-1 in HEK293T cells treated with PBA for a series of different time duration over treated with DMSO alone for 24 hrs. GABA uptake activity was measured by the high-throughput ^3^H radio-labeling GABA uptake on a liquid scintillator with QuantaSmart. **C.** Cartoon showing the mutant allele only was compared. HEK293T cells were transfected with the wildtype or the mutant GAT-1^YFP^ cDNAs alone for 48 hr. **D**. Cartoon showing the coexistence condition of the wildtype and the mutant allele in patients. HEK293T cells were transfected with the wildtype GAT-1^YFP^ alone or in mixture of the wildtype or the mutant cDNAs for 48 hr. In the mixed condition, the ratio of the wildtype GAT-1 with pcDNA or the mutant cDNAs are 1:1 with the total cDNA amount of 0.5 μg. PBA (2mM) was applied for 24 hrs while DMSO was applied as control. Both wt and the mutant were normalized to its own DMSO treated conditions. 966 stands for the wildtype treated with Cl-966 (50 um) while 711 stands for NNC-711 (35um). (N=4-7 different transfections. In **A** and **B**, *P<0.05, **P<0.01, \*\*\**p*< 0.001 vs wt 0, In **C** and **D**, *P<0.05, **P<0.01, \*\*\**p*< 0.001 vs its own DMSO treated. In **C**, ns stands for no significance. Unpaired t test or one-way analysis of variance (ANOVA) and Newman-Keuls test. Values were expressed as mean ± S.E.M.)

Because all patients carrying *SLC6A1* variants are heterozygous and only one allele is affected in patients as illustrated (Figure 3C), we thus tested the effect of PBA in either “homozygous” with the mutant cDNAs alone or “heterozygous” with a mixture of the wildtype and the mutant cDNAs. In the mutant alone, PBA did not increase GABA uptake for the GAT-1(E16X) and GAT-1(L460R) mutations (0.813±0.06 for E16X, 1.378±0.056 for V125M, 1.22±0.063 for A288V, 1.386±0.058 for S295L, 1.185±0.065 for G362R, 1.236±0.074 for D410E, 1.14 ± 0.07 for L460R and 1.323± 0.054 for W495X) (Figure 3C). However, PBA increased the mutant GAT-1 activity in all tested variants except the GAT-1(E16X) in the “heterozygous” condition, which reflects the patient condition in which a wildtype allele is present (1.149 ±0.025 for E16X, 1.33±0.05 for V125M, 1.248±0.053 for A288V, 1.318±0.062 for S295L, 1.262±0.026 for G362R, 1.17±0.035 for D410E, 1.214 ± 0.037 for L460R and 1.305± 0.0296 for W495X) (Figure 3D). The failure to increase GABA activity in GAT-1(E16X) when expressed alone is most likely because of the very short peptide resulting from the early truncation, rendering it very misfolded and lacking ability to be refolded, thus being subjected to quick degradation inside the ER. In the “heterozygous” condition, in which the cells transfected with mixed wildtype and mutant GAT-1 cDNAs, all the mutant conditions had increased GABA uptake activity compared with the DMSO treated conditions.

### PBA increased GABA uptake in the patient iPSC derived astrocytes

We then determined if PBA could restore the GAT-1 activity in human patient iPSC-derived astrocytes. We use astrocytes because GAT-1 is expressed in both astrocytes and neurons but there is a direct correlation of astrocytic GAT-1 deficit with thalamic absence seizures^27,28^. We first differentiated the iPSCs to neuronal progenitor cells (NPCs) and then differentiated the NPCs into astrocytes. Human astrocytes derived from iPSCs at days 30 to 35 after differentiation were treated either with DMSO or PBA (2mM) for 24 hrs before GABA uptake assay. Based on our protocol, > 90% of astrocytes adopts typical star-like astrocytic morphology after day 27 of differentiation (Figure 4A). When compared with the isogenic control cell line, the patient astrocytes had reduced GABA uptake activity as we previously reported^11^. However, PBA (2mM) treatment for 24 hrs increased the activity in both isogenic cells and patient cells and increased the uptake of the patient cells from 57% to 81% of the corrected cells (Figure 4B). The increased magnitude is larger in the mutant than the isogenic control astrocytes (1.173 ± 0.026 for control vs 1.387±0.08 for patient) (Figure 4C). We then tested the effect of PBA on all eight mutations in this study in human astrocytes. We transfected the wildtype or the mutant GAT-1^YFP^ (1μg cDNAs/per 35 mm^2^ dishes) in the astrocytes derived from the corrected iPSCs for 48 hrs. GABA uptake activity was determined in astrocytes treated with DMSO or PBA 2mM for 24 hrs. GAT-1 inhibitor Cl-966 (50μM) or NNC-711 (35 μM) was applied to make sure the specificity of the detected radioactive signal. PBA increased GABA activity in the wildtype and all the mutant conditions (1.188±0.075 for wt; 1.182±0.025, E16X, 1.285±0.018 for V125M, 1.204±0.031 for A288V, 1.353±0.065 for S295L, 1.25±0.023 for G362R, 1.21±0.022 for D410E, 1.18 ± 0.045 % for L460R and 1.45± 0.04 for W495X). However, the GABA activity was increased the most in the astrocytes expressing the mutant GAT-1(W495X) (Figure 4D).

**Figure 4.**
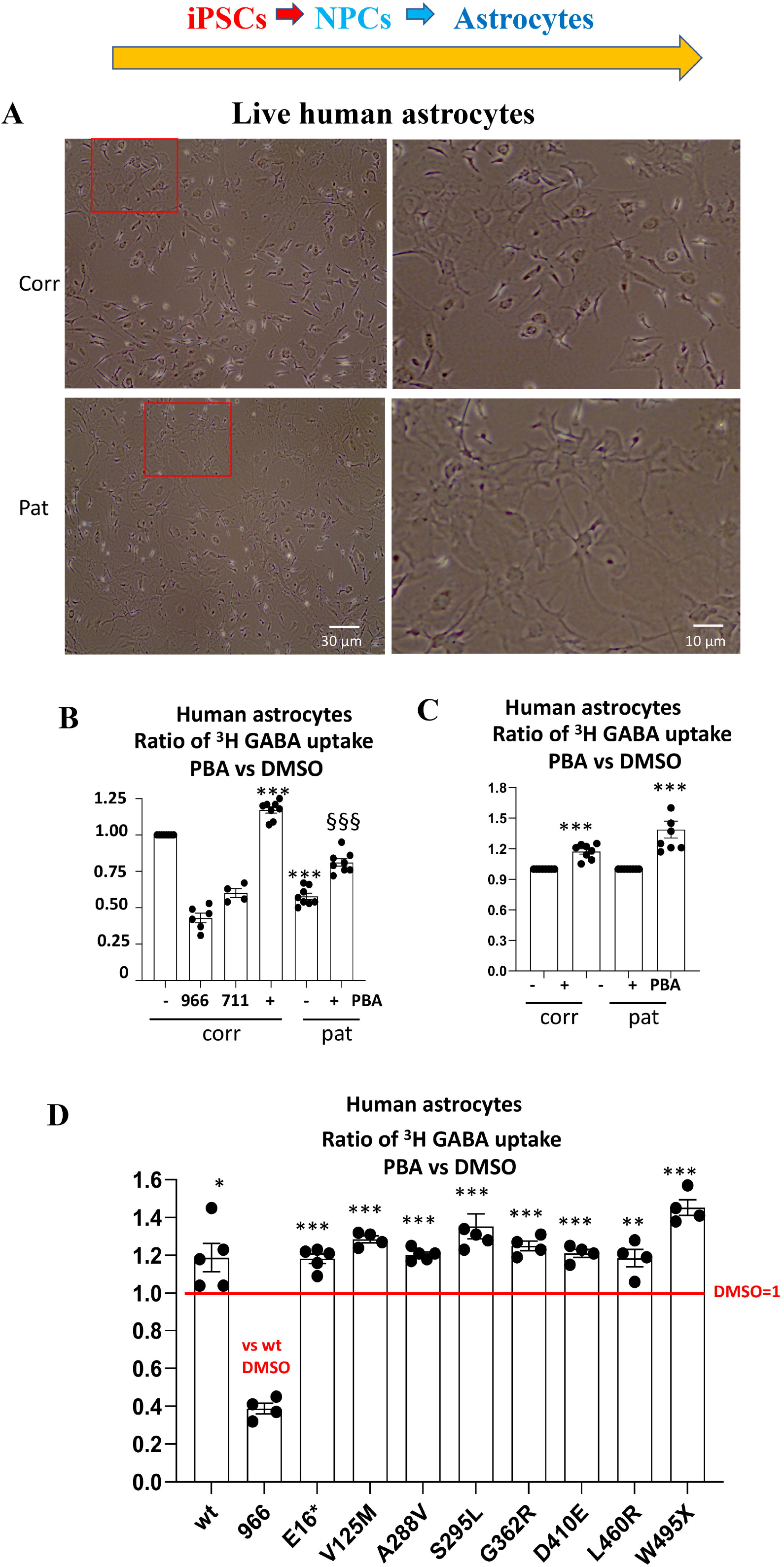
4-Phenylbutyrate rescued the GABA uptake function in the patient astrocytes and in human iPSC derived astrocytes over-expressing the mutant GAT-1 transporters. **A.** Live images of human astrocytes differentiated from the patient (pat) or its CRISPR corrected isogenic control (corr) iPSCs at day 30 (**A**). **B, C.** Astrocytes at day 30 to 35 after differentiation were treated with PBA (2mM) for 24 hrs before ^3^H radioactive GABA uptake assay. The DMSO treated corr or pat was taken as 1. **D.** Corrected astrocytes at day 30-35 after differentiation were transfected with the wildtype or the mutant GAT-1^YFP^ cDNAs (1μg per a 35mm^2^dish) for 48 hrs before ^3^H radioactive GABA uptake assay. PBA (2mM) was applied for 24 hrs before GABA uptake assay experiment. GABA flux was measured after 30 min transport at room temperature. The influx of GABA, expressed in pmol/μg protein/min, was averaged from duplicates for each condition and for each transfection. The average counting was DMSO treated condition taken as n = 100%. In **B** and **D**, 966 stands for Cl-966 (50μm) and &11 stands for NNC-711 (35μm) that was applied 30 min to the astrocytes transfected with the wildtype GAT-1^YFP^ before preincubation and removed during preincubation. In **B,** ***p < 0.001 vs. corrected DMSO treated. §§§p<0.001 vs patient DMSO treated. In **C,** the PBA treated corrected or patient GABA uptake was normalized to its DMSO treated. In **D,** the PBA treated wildtype or mutant GABA uptake was normalized to its DMSO treated. *p < 0.05; **p < 0.01; ***p < 0.001 vs. DMSO treated in its own group. In **B** and **C**, n=4-8 batches of cells. In **D**, n=4-5 different transfections). Unpaired t test. Values were expressed as mean ± S.E.M).

### The human iPSC derived astrocytes expressing mutant GAT-1 (S295L) caused retention of the wildtype GAT-1 inside the ER

We have previously demonstrated that GAT-1(S295L) was retained inside ER^11^. However, it is unknown if the mutant GAT-1 would suppress the wildtype GAT-1 via aberrant oligomerization of the wildtype GAT-1 with the mutant GAT-1. We then compared the GAT-1 expression pattern in the corrected or the patient iPSCs derived astrocytes. Human astrocytes derived from iPSCs at days 30 to 35 after differentiation were cotransfected with the wildtype or the mutant GAT-1 cDNAs with the ER marker ER^CFP^ (Figure 5A). To evaluate the subcellular localization of the mutant GAT-1, we measured the colocalization fluorescence of the GAT-1-representing YFP and ER-representing CFP in the corrected or the patient astrocytes coexpressing the wildtype GAT-1^YFP^ or the mutant GAT-1^YFP^ with ER^CFP^. It is not surprising that the mutant GAT-1(S295L) had increased overlapping fluorescence signal with ER than the wildtype. Importantly, the wildtype GAT-1 expressed in the patient astrocytes had increased ER overlapping fluorescence signal than in the corrected cells (36.29±2.4% for GAT-1^YFP in^ corrected cells, 56.6±1.95% for GAT-1^YFP^ in patient cells, 77.78±3.26 % for GAT-1 (S295L)^YFP^ in corrected cells, 92.49±0.99% for GAT-1 (S295L)^YFP^ in patient cells (Figure 5B). We then determined if PBA could increase the GAT-1 protein expression in the human astrocytes and found PBA treatment increased the GAT-1 expression in both the corrected and the patient conditions (1.228± 0.04 for correct vs 1.41 ±0.083 for patient) (Figure 5C, D, E). The total GAT-1 protein of the patient astrocytes was increased from 60.8% to 84.6% of the corrected. We could not determine the surface GAT-1 expression because of the low yield of protein in astrocytes. It is likely that the increased GAT-1 in the patient astrocytes is contributed by the wildtype GAT-1. More detailed characterization with specific tag in GAT-1 to distinguish the wildtype vs the mutant allele may further elucidate the contribution of each allele.

**Figure 5.**
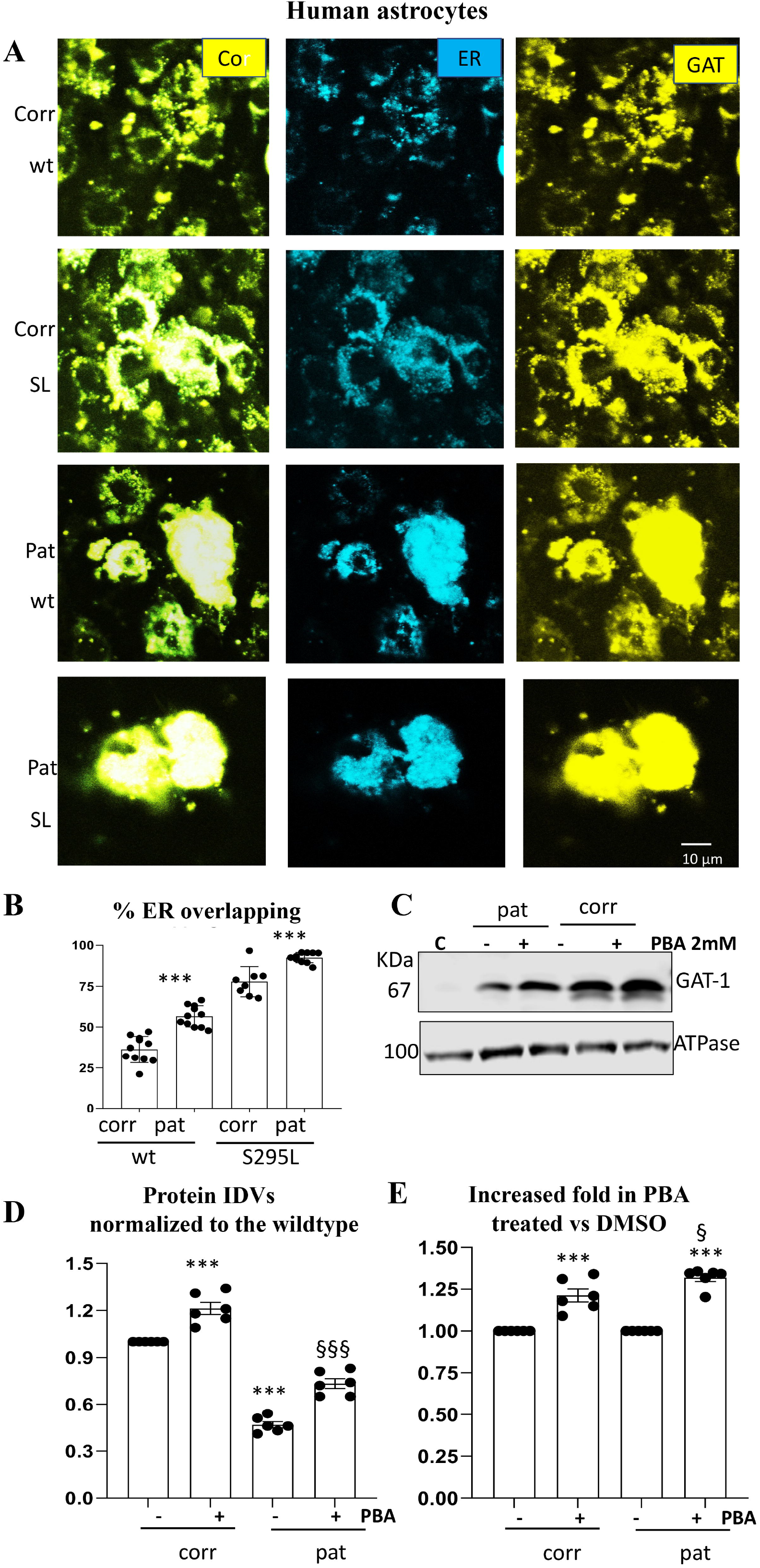
The patient astrocytes caused ER retention of the wildtype and exacerbate the mutant GAT-1 ER retention while 4-Phenylbutyrate increased GAT-1 protein expression. **A-B.** Human patient corrected (isogenic control, Corr) or uncorrected patient (Pat) astrocytes at day 30 to 35 after differentiation from iPSCs were co-transfected with the endoplasmic reticulum (ER) marker ER^CFP^ in combination with the wildtype or the mutant GAT-1^YFP^ cDNAs (0.5μg:0.5 μg per a 35mm^2^dish) for 48 hrs before confocal microscopy analysis. Confocal images were acquired in live astrocytes under 63X objective with zoom under 2.5 (**A**). **B**. The graph represents the ER overlapping signal of GAT-1^YFP^ analyzed by Metamorph. **C**. **D.** Total cell lysates from astrocytes cultured in 100 mm^2^ dishes treated with DMSO or PBA (2mM) for 24 hrs and were analyzed with SDS-PAGE. Membranes were blotted with a rabbit polyclonal atnti-GAT-1 antibody (**C**). The lysates of CHO cells were used as control. **D, E**. The protein IDVs of the corrected or patient GAT-1 and in human astrocytes was normalized to the corrected astrocytes treated with DMSO, the GAT-1 protein was normalized to its own internal control ATPase or GAPDH and then to the DMSO treated corrected which is arbitrarily taken as 1 (**D**) or its own DMSO treated is taken as 1 (**E**). Values were expressed as mean ± S.E.M). In **B**, N=8-11 fields from 4 different transfections in 4 different batch of astrocytes. In **C**, **D, E**. N=6 batch of cells differentiated from 4 different batches of neural progenitor cells, In **B**, \*\*\**p*< 0.001 vs corrected. In **D**, *** P<0.001 corrected untreated. zzzP<0.001 vs Patient untreated. In **E**, \*\*\**p*< 0.001 vs its own untreated; zP<0.05 vs corrected PBA treated, one-way analysis of variance and unpaired t test).

### PBA increased GABA uptake in the astrocytes from mutationbearing knockin mice

We then determined the effect of PBA in astrocytes from knockin mice. We chose to investigate the GAT-1(A288V) and GAT-1(S295L) because we have extensively studied these two mutations^11^. We used cortical astrocytes because of the cortico-thalamic pathway involved in absence seizures. Additionally, we found the data from the cortical astrocytes and thalamic astrocytes to be comparable (data not shown). We used cortical astrocytes because the tissue is more abundant than thalamus. Because GABA uptake activity can be contributed to by both GAT-1 and GAT-3, we treated the cells with GAT-3 inhibitor SNAP5114 (30 μM) to ensure only GAT-1 activity was measured. Compared with the wildtype, the astrocytes cultured from the heterozygous pups had reduced GABA uptake activity (0.38±0.021 for A288V and 0.60±0.06% for S295L) (Figure 6A). PBA treatment increased the GABA uptake activity in the astrocytes cultured from the wildtype and the mutant pups (1.22 ±0.058 for wt; 1.26 ±0.063 for A288V and 1.54±0.13 for S295L) (Figure 6B). The magnitude of increase in the astrocytes cultured from *Slc6a1^+/S295L^* pups tended to be larger than that from the wildtype though all showed an increase in GABA uptake.

**Figure 6.**
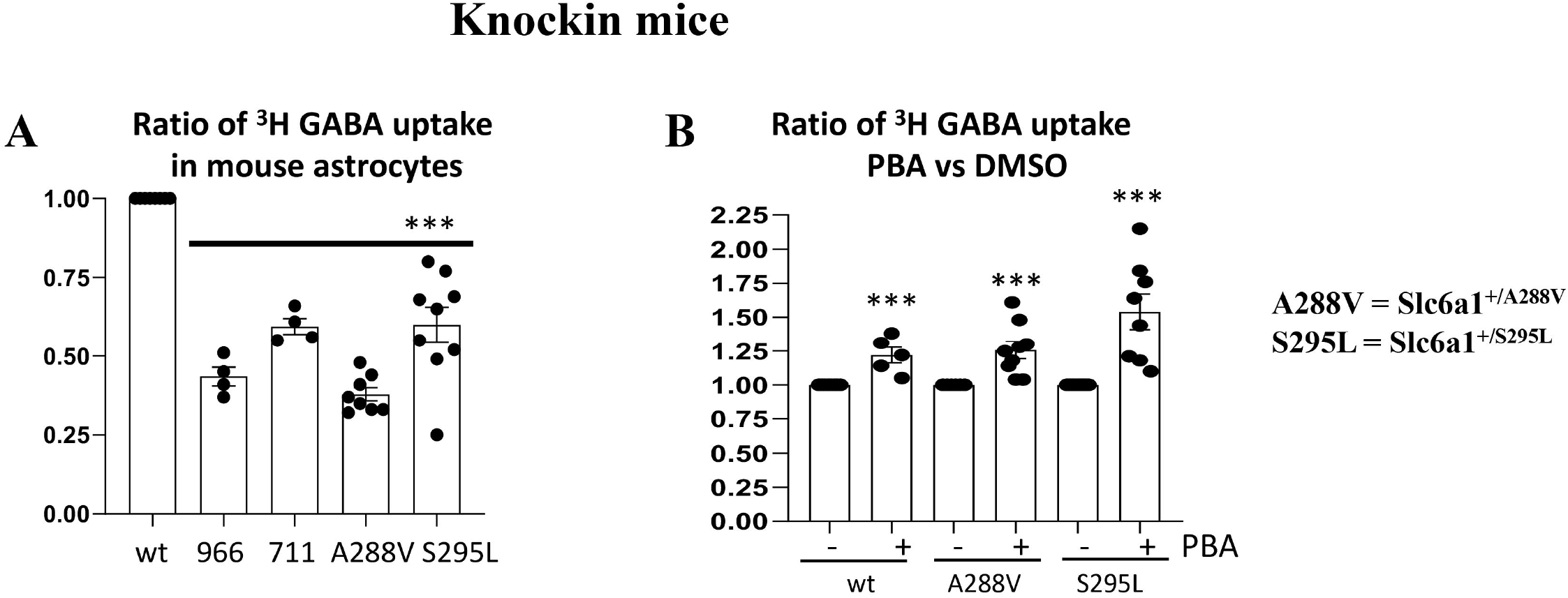
4-Phenylbutyrate rescued the GABA reuptake function in cortical astrocytes cultured from two knockin mouse lines carrying *Slc6al* mutations. Mouse cortical astrocytes were cultured from postnatal day 0-3 days old pups from the Slc6a1^+/A288V^ or Slc6a1^+/S295L^ mice. **A**. Astrocytes under passage 2 were grown in 100-mm^2^ dishes and passaged into 35-mm^2^ dishes before GABA uptake assay. Cl-966 (100μm) and NNC711 (70μm) was applied 30 min before preincubation and removed during preincubation. GAT-3 inhibitor (SNAP5114 (30μM) was applied during GABA uptake to make sure only GAT-1 activity was measured. The wildtype of either Slc6a1^+/A288V^ or Slc6a1^+/S295L^ were taken as 1. The cultures derived from each mutant mouse was compared to the culture from its own wildtype littermates. **B**. Sister cultures of astrocytes from different mouse lines were applied with DMSO or PBA 2mM for 24 hr before GABA uptake. The wildtype data were pooled from two mouse lines. The GABA uptake activity of PBA treated was normalized to its own DMSO conditions. The graph represents the relative GABA uptake level normalized to cells of its own genotype treated with DMSO. (***p < 0.01 vs. wt, n=4-9 batch of astrocytes from 4 pairs of littermates for **A** and 5-8 batches of astrocytes from 4 pairs of littermates-9 for **B**). Cl-966 (100μm) was applied 30 min before preincubation and removed during preincubation One-sample t test. Values were expressed as mean ± S.E.M).

### GAT-1 protein expression was reduced in *Slc6a1^+/A288V^* and *Slc6a1^+/S295L^* knockin mice, which was partially restored by PBA

The increased GABA uptake activity could be due to increased GAT-1 protein expression caused by promoted membrane trafficking by PBA. We then determined if PBA alters the GAT-1 in knockin mice. We first profiled the GAT-1 protein expression of the mutant mice. We then treated the mice between 2-4 months old with vehicle or PBA for 1 week. The lysates from cortex, cerebellum, hippocampus, and thalamus were surveyed. Compared with the wildtype, the heterozygous mice from both the *Slc6a1^+/A288V^* and *Slc6a1^+/S295L^* mouse lines had reduced GAT-1 expression in all surveyed brain regions (wt: 1.025 ±0.023 for cortex, 0.86 ±0.03 for cerebellum; 1.07 ±0.03 for hippocampus; 1.22 ±0.033 for thalamus; *Slc61^+/A288V^* het: 0.57 ±0.0165 for cortex; 0.51 ±0.056 for cerebellum; 0.547 ±0.029 for hippocampus; 0.58 ±0.037 for thalamus; *Slc61^+/S295L^* het: 0.513 ±0.023 for cortex; 0.465 ±0.021 for cerebellum; 0.55 ±0.023 for hippocampus; 0.50 ±0.016 for thalamus), consistent with previous findings *in vitro* that the GAT-1(A288V) and GAT-1(S295L) mutations cause ER retention of the mutant protein, consequently leading to enhanced degradation^11^. Compared with vehicle treated littermates, PBA treatment increased GAT-1 expression in the cortex, hippocampus, and thalamus after normalization with the housekeeping ATPase. PBA increased the total expression of GAT-1 in cortex, hippocampus, and thalamus but this was not observed in cerebellum (*Slc61^+/A288V^* PBA: 1.29 ±0.05 for cortex, 1.045±0.028 for cerebellum; 1.28 ±0.044 for hippocampus; 1.35 ±0.049 for thalamus; *Slc61^+/S295L^* PBA: 1.168 ±0.033 for cortex; 1.03 ±0.03 for cerebellum; 1.30 ±0.04 for hippocampus; 1.26 ±0.04 for thalamus vs the same brain region of the vehicle treated mice taken as 1). This may suggest that the PBA-induced increase of GAT-1 can be region-specific and that the increased membrane trafficking of GAT-1 contributes to increased GABA uptake. It is worth noting that the PBA treatment did not increase GABAA receptor γ2 subunit (Supplementary Figure 4), suggesting that the PBA-induced protein increase could be mutation-bearing gene specific, but this merits more elucidation.

### PBA reduced seizure activity in *Slc6a1^+/S295L^* knockin mice

A major phenotype in *SLC6A1* mutation mediated disorders is epilepsy, leading us to investigate whether PBA reduces seizure activity in vivo. We tested the effect of PBA in *Slc6a1^+/S295L^* mice. First, 2–3-month-old mice were implanted with EEG head mounts. After 5-7 days recovery, the mice were treated with a normal saline vehicle, and then subject to video monitored EEG recording for 48 hrs. After at least one day rest, the mice were administered with PBA (100mg/kg) for 7 days followed by 48-hour EEG recording (Figure 8A). We observed frequent absence-like seizures and occasionally generalized tonic clonic seizures during routine handling in *Slc6a1^+/S295L^* mice. In EEG recordings, the *Slc6a1^+/S295L^* mice displayed frequent absence like activity (5-7 Hz) and myoclonic jerks with or without behavioral correlation (Figure 8B and C). However, compared with vehicle treated baseline recordings, PBA treatment reduced the absence seizure representing 5-7 Hz SWDs from 149.5 with vehicle treated to 45.5 over 48 hrs. of recordings (Figure 8C, D). Importantly, every mouse had reduced seizures compared to vehicle treatment ranging from 55% to 89% with 11% to 44.6% seizure activity remaining (Figure E, F). This suggests PBA alone reduced seizure activity in *Slc6a1^+/S295L^* mice. This is likely due to promoted membrane protein trafficking resulting from increased functional GAT-1 expression. It is worth noting that the increase of GAT-1 is specific. There was no increase of the expression of GABAA receptor subunit such as γ2 subunit (Supplementary Figure 4).

**Figure 7.**
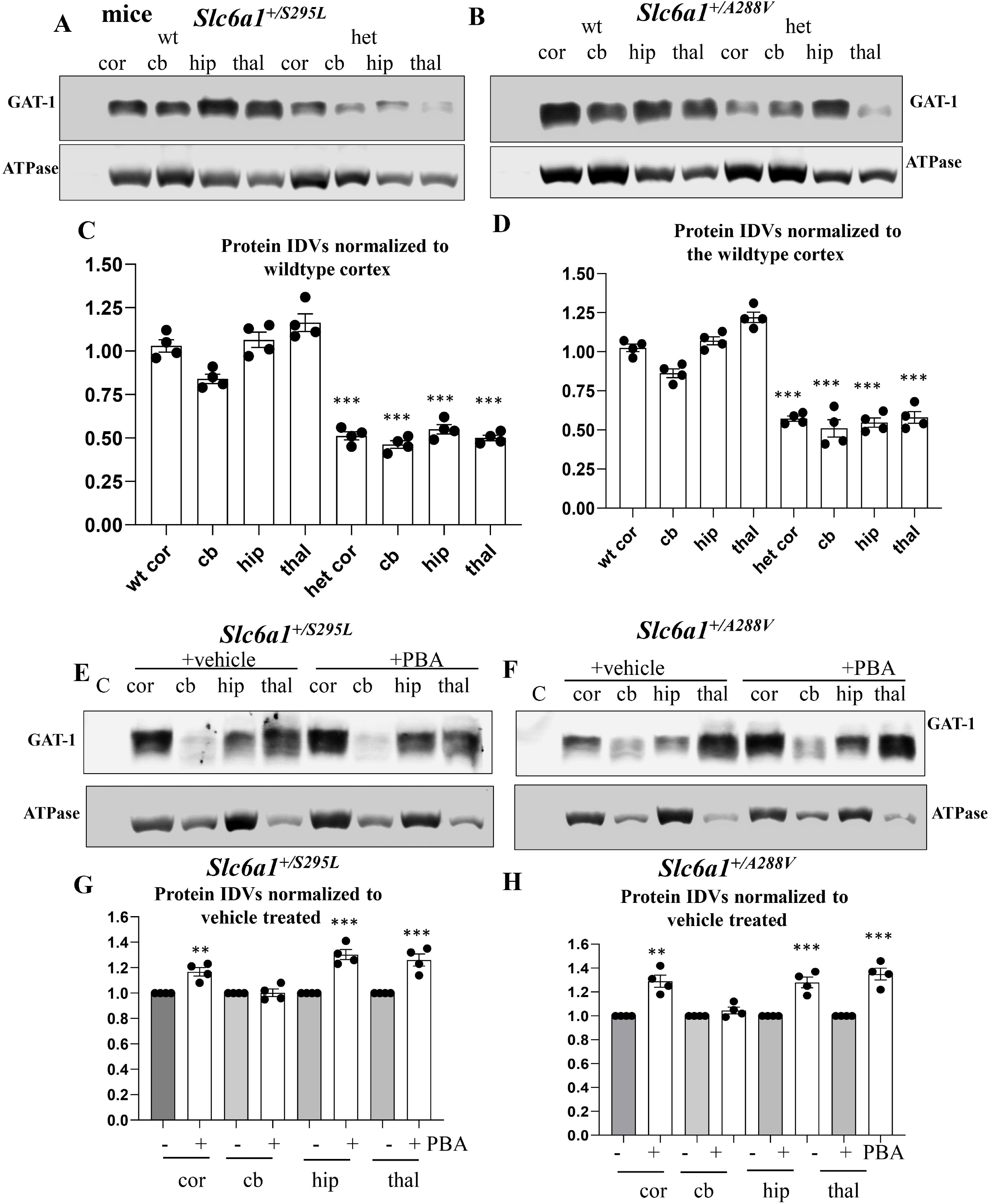
Both A288V and S295L mutation bearing knockin mice had reduced GAT-1 protein that was partially restored by 4-Phenylbutyrate treatment. **A-H.** Lysates from different brain regions (cortex (cor), cerebellum (cb), hippocampus (hip) and thal (thalamus) from the wildtype (wt) and heterozygous (het) mice at 4-6 months old, untreated (**A, B**) or treated with vehicle or PBA (100mg/kg) for 7 days (**E, F**) were subjected to SDS-PAGE and immunoblotted with anti-GAT-1 antibody. (**C**, **D**). Integrated density values (IDVs) for total GAT-1 from wild-type and het KI were normalized to the Na^+^/K^+^ ATPase or anti-glyceraldehyde-3-phosphate dehydrogenase (GAPDH) loading control (LC) in each specific brain region and plotted. N=4 from 4 pairs of mice. (**G, H**). Integrated density values (IDVs) for total GAT-1 from het KI treated with vehicle or treated with PBA were normalized to the Na^+^/K^+^ ATPase or anti-glyceraldehyde-3-phosphate dehydrogenase (GAPDH) loading control (LC). The IDVs of the heterozygous treated with PBA was then normalized to vehicle treated. The vehicle treated in each brain region was taken as 1. N=4 from 4 pairs of mice for **C, D, G** and **H**, Values were expressed as mean ± S.E.M. One way ANOVA or unpaired t test. In C and D, ***p < 0.001 vs. wt, in G and H, **p < 0.01; ***p < 0.001 vs vehicle treated).

**Figure 8.**
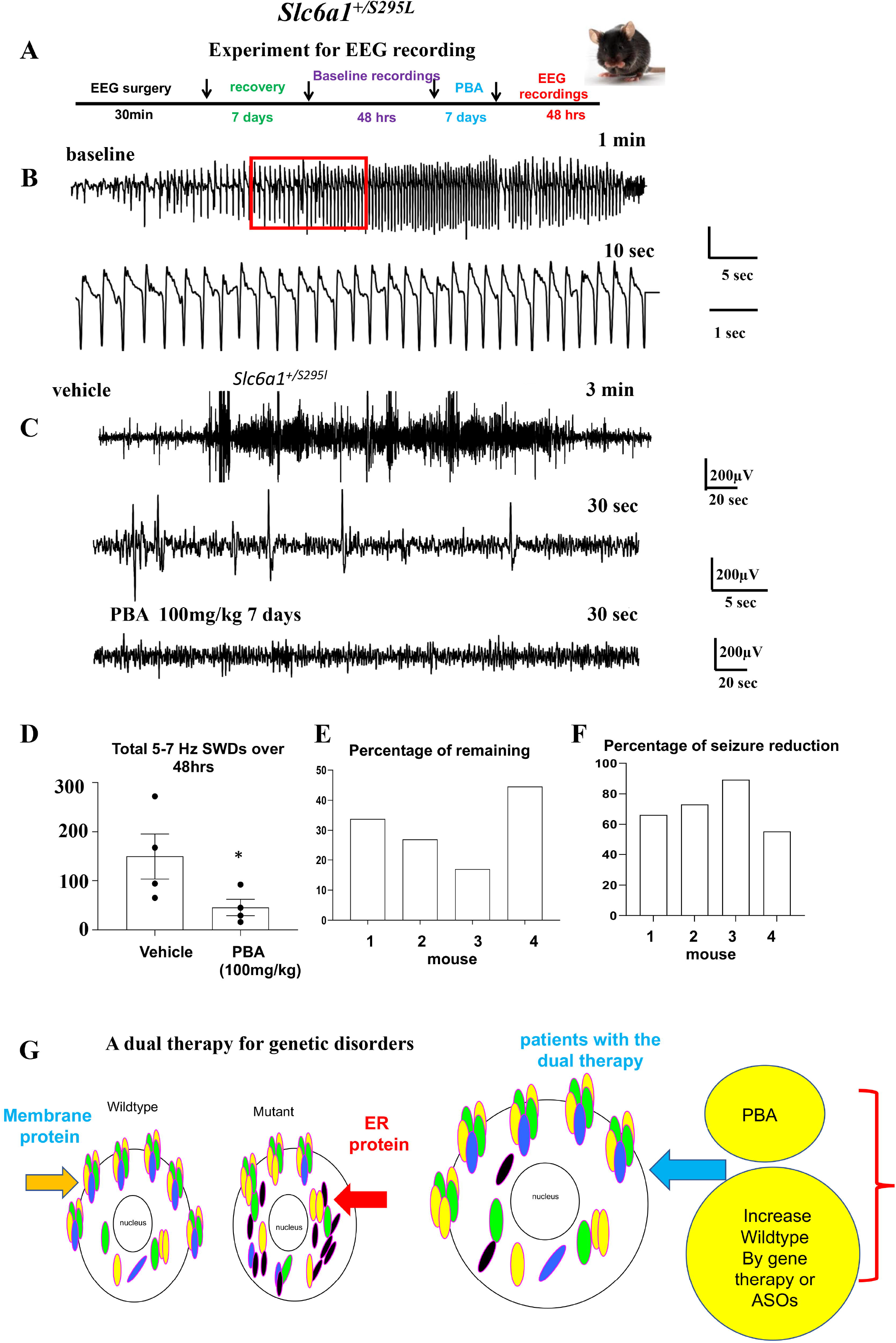
4-Phenylbutyrate alone reduced seizures in *SLc6a1^+/S295L^* knockin mouse. **A**. Schematic depiction of experimental paradigm for EEG recordings and PBA treatment. **B**. Representative EEG recordings show that the heterozygous *Slc6a1^+/S295L^* (het) KI mice had frequent absence like spike wave discharges (SWDs) and myoclonic jerks during baseline recordings. **C**. Comparison of EEG traces recorded after the vehicle (normal saline 100 μl) treated or after treatment with PBA (100mg/kg, ip, single dose, daily) for 7 days. **D.** Graph showing the total number of 5-7 Hz SWDs calculated by Seizure Pro during 48 hrs recordings after vehicle or PBA treatment. PBA treatment (100mg/kg, ip, single dose, daily) for 7 days reduced seizure activity. *p < 0.05; vs vehicle treated, paired t test, N=4, Values were expressed as mean ± S.E.M). **E, F.** Graph showing the percentage of seizure remaining (**E**) and seizure reduction (**F**) in each mouse after PBA treatment (N=5 mice, paired t test). **G.** Based on our findings, we propose a dual therapy as a feasible approach for treating *SLC6A1* mutations and many other genetic disorders by boosting the wildtype allele, removing the mutant allele and ER stress with PBA.

## Discussion

### Reduced membrane trafficking due to ER retention is a major cause of *SLC6A1* pathological variants

*SLC6A1* variants are associated with a wide spectrum of neurodevelopmental disorders ^9,13,31^. We have previously studied the impact of the variants on GABA uptake function, membrane trafficking and subcellular localizations of the mutant protein in both neurons and astrocytes in mouse and human iPSC derived GABAergic neurons and astrocytes^11^. We found that the mutant GAT-1 stemming from missense mutations resulted in complete or partial loss of GABA uptake function while the premature codon generating nonsense mutations resulted in nearly complete loss of function^11–13^. The mutant protein is often retained inside the ER and subject to enhanced degradation as observed in many GABA_A_ receptor subunit mutations, suggesting that *SLC6A1* mutation caused disorders could be rescued with a common treatment despite the highly heterogenous clinical phenotype.

### The impaired membrane trafficking resulted in reduced functional and mature GAT-1 but increased nonfunctional, immature GAT-1

What are the molecular mechanisms underlying the reduced surface expression of the mutant GAT-1? For a membrane protein such as GAT-1 or GABA_A_ receptor subunits, only those protein that are correctly folded and fully glycosylated can traffic to the cell surface or synapse where it exerts biological function. The protein that is inside the ER has no biological function. By contrast, the ER retained mutant protein would prevent the wildtype protein from correct oligomerization and trafficking, and cause ER stress in the cell. Endo-H digestion can distinguish the mature form of GAT-1 vs immature GAT-1 because the ER-attached glycan is sensitive to Endo-H digestion. Based on our findings, the mutant GAT-1 resulted in decreased functional mature GAT-1 but increased amount of immature ER-retained GAT-1 except the GAT-1(E16X) that is likely subject to rapid disposal. The similar ER retention of the GAT-1 was reported in astrocytes in our previous studies^11^. Therapeutic strategies that increase the mature GAT-1 and reduce the immature GAT-1 should be beneficial for many mutations.

### The mutant GAT-1 can prevent forward trafficking of wildtype GAT-1, thus reducing the function of the wildtype allele

The oligomerization status and detailed structure of membrane bound GAT-1 remain unknown. It is possible that wildtype GAT-1 and mutant GAT-1 can form dimers or other high molecular mass protein complexes and the trafficking deficient mutant GAT-1 could potentially interfere with the trafficking of the wildtype. This notion is evidenced in the human astrocytes, where wildtype GAT-1 was more ER bound when expressed in the patient cells than expressed in the corrected cells (Figure 5). The GAT-1(S295L) protein is retained inside the ER with minimal surface expression. In the patient cells expressing the mutant GAT-1(S295L)^YFP^, the GAT-1-representing yellow fluorescence formed large clumps inside cells with enlarged ER, suggesting substantial ER retention of the mutant protein inside the ER. PBA increased GABA uptake activity and the total GAT-1 protein in both the corrected and patient astrocytes.

### PBA could rescue the GABA uptake activity across cell types

We tested the effect of PBA in heterologous cells, mouse, and human astrocytes and found that PBA increased GABA uptake in different cell types. It is worth noting that we observed the increase of GABA uptake in neural progenitor cells after PBA treatment, although the baseline activity of GABA uptake in NPCs is relatively low. This is consistent with our previous study on comparison of GABA uptake in NPCs, astrocytes, and inhibitory neurons^11^. This suggests that the protein quality control machinery is conserved across species and cell types. This is important considering the very early onset of the *SLC6A1* variant-mediated disorders in human patients and the expression of GAT-1 function in multiple cell types. This indicates PBA treatment can increase GAT-1 function before and after the postmitotic mature neurons are formed and that PBA can improve the function of GAT-1 in both progenitor cells and the derived neurons and astrocytes. This indicates that PBA treatment is likely disease modifying because it can improve GAT-1 function in all involved cell types instead of simply masking disease symptoms.

### PBA increased the GAT-1 activity in the heterozygous condition likely through the wildtype allele and in some case, the functional mutant allele

PBA is a hydrophobic chaperone, and it may prevent the aberrant interaction of the mutant GAT-1 protein with its wildtype binding partners. For the mutant allele alone, PBA increased most mutant GAT-1 but not E16X. This is likely because the GAT-1(E16X) is severely misfolded, with only 15 amino acids left in the protein peptide and could not be rescued. PBA increased GABA uptake for all surveyed mutations in the heterozygous conditions when the wildtype allele is present. The increase of function in the heterozygous condition is likely mainly contributed by the wildtype allele. In some cases, such as GAT-1(A288V), the mutant allele could also be functional if rescued and present on the cell surface. This is critical, since all *SLC6A1* mutation-bearing patients identified so far are heterozygous, suggesting that they can potentially benefit from a treatment option like PBA. Based on our data, it is likely that the effect of PBA is bidirectional. For the wildtype allele, it can promote membrane trafficking while facilitating the disposal of those severely impaired proteins, thus making the trafficking more conducive for the wildtype allele. Because the patients are heterozygous, this will consequently increase the net GABA uptake in patient conditions. In fact, in patient iPSCs derived cells, PBA increased GAT-1 expression in both the corrected and patient conditions. In mutation knockin mice, PBA increased GAT-1 expression in cortex, hippocampus, and thalamus, implicating a relatively less disturbed cortico-thalamic-cortical circuitry.

### PBA treatment increased functional GAT-1 and reduced seizures in mutation knockin mice

Seizures and abnormal EEGs are common among patients with *SLC6A1* mutations, and thus can serve as a good biomarker for evaluating the effect of PBA. GAT-1(S295L) protein alone has no GABA function as described in here and in our previous work^11^. In *Slc6a1^+/S295L^* mice, we found significant loss of total GAT-1 in all major brain regions. PBA treatment for 1-week reduced seizures in the *Slc6a1^+/S295L^* heterozygous mice, likely via increasing the functional GAT-1. The magnitude of GAT-1 increase is variable in different brain regions. The magnitude of increase of GABA uptake and GAT-1 protein expression is modest. Based on our studies on other epilepsy mouse models *Gabrg2^+/−^* and *Gabrg2^+/Q390X^*, a small increase of γ2 subunit reduces seizure severity from Dravet syndrome to seizure free^35^. This is likely true for the PBA treated condition, and the increase of functional GAT-1 reduced the seizure burden in *Slc6a1^+/S295L^* mice by ~70% with PBA treatment alone. PBA increases GABA uptake activity. Deficits in GABA uptake is a common mechanism underlying the heterogenous clinical phenotypes associated with SLC6A1 mutations. This suggests that PBA could be applied to patients with the same molecular defects regardless of clinical phenotype, including various epilepsy syndromes, autism, neurodevelopmental delay among others.

### A feasible dual therapy for *SLC6A1* mutations and possibly for many other genetic disorders

There is no effective treatment for *SLC6A1* mutation mediated disorders to date. Valproic acid has been reported to control seizures but could not improve cognition^13^. This suggests that the impaired cognition is likely caused by the deficit of GAT-1 function instead of being secondary to seizure activity. Although the detailed mechanisms of action for PBA needs more thorough investigation, we believe the prompted membrane trafficking and increased functional GAT-1 is a major mechanism underlying the seizure mitigation. There are numerous ion channels and transporters associated with epilepsy, autism, and neurodevelopmental delay. Based on our substantial characterizations of the impact of mutations in GABA_A_ receptors and more recently in GAT-1, we propose that impaired trafficking and ER-associated degradation (ERAD) are common mechanisms for mutations in GABA receptors, transporters, and beyond. This study, in combination with our previous work on the patho-mechanisms of GABA receptors and transporter 1 mutations, provides evidence that PBA may be a feasible treatment option and is disease-modifying. Considering PBA is FDA-approved for pediatric use and is orally bioavailable and that gene therapy and antisense oligonucleotides (ASOs) are on the horizon, this treatment could open a new door or even bring a cure for many genetic disorders by boosting the wildtype functional allele via gene therapy and reducing the ER-retained mutant protein and ER stress via PBA as illustrated in Figure 8G. We propose this dual therapy is an actionable mode of treatment for *SLC6A1* variants mediated disorders and far beyond.

(Note: our work has prompted a pilot clinical trial and the feedback from patients is very promising. The parents of the three patients S295L, G362R and a *SLC6A1* deletion patients all reported improved overall quality of life, including reduced seizures, reduced autistic features and better cognition).

## Ethics statement

All animals and related experiments in this study were approved by the Vanderbilt University IACUC.

## Acknowledgements

The work of functional evaluation was carried out at Vanderbilt University Medical Center and was supported by research grants from National Institute of Health (NINDS) NS82635 to KJQ. The work was also supported by research grants from *SLC6A1* Connect and Taysha Gene Therapies to KJQ). The patient derived and CRISPR corrected induced pluripotent stem cells were kindly shared by Drs. Jason Aoto and Scott Demarest in University of Colorado). Imaging data were performed in part through the VUMC Cell Imaging Shared Resource. *correspondence).

## Author contributions

KJQ conceived the project. FM and KJQ performed radiolabeling GABA uptake. FM, GN and KJQ performed flow cytometry. KR and KJQ performed iPSC derived cell cultures. SP did transfection and astrocyte. WS and SP performed biochemistry experiments and protein assays for radiolabeling GABA uptake assay. GN performed drug administration and EEG recordings in mice. CF did EEG scoring. KJQ and GN wrote the paper. All authors reviewed and edited the compiled manuscript.

## Supplementary material

Supplementary material is available at *Brain* online.

---

Supplementary Figure 1: full-length gels of immunoblots Figure 2A and Figure 2B. Supplementary Figure 2: Original full-length gels for Figure 5C.

Supplementary Figure 3: Original full-length gels for Figure 7A, 7B, 7E and 7F.

**Supplementary Figure 4: The GABA_A_ receptor γ2 subunit protein was not increased in 4-phenylbutyrate treated *Slc6a1^+/S295L^* mice. A, B. Original full-length gels for The GABA_A_ receptor γ2 subunit protein in different brain regions of the heterozygous mice treated with vehicle or PBA. C. Graph showing the fold change of protein IDVs from mice treated with PBA normalized to its loading control ATPase and then to the vehicle treated.** The heterozygous (het) mice at 2-8 months old were treated with vehicle or PBA (100mg/kg) for 7 days. The total lysates from cortex (cor), cerebellum (cb), hippocampus (hip) and thalamus (thal) were subjected to SDS-PAGE and immunoblotted with rabbit anti-γ2 antibody. Integrated density values (IDVs) of the γ2 subunit from het mice treated with PBA were normalized to the Na^+^/K^+^ ATPase or anti-glyceraldehyde-3-phosphate dehydrogenase (GAPDH) loading control (LC) and then to the vehicle treated, which is arbitrarily taken as 1 in each specific brain region. N=4 from 4 pairs of mice.

