## Supplementary figures and images for "4-phenylbutyrate restored GABA uptake and reduced seizures in *SLC6A1* variants-mediated disorders"

### Supplementary 1

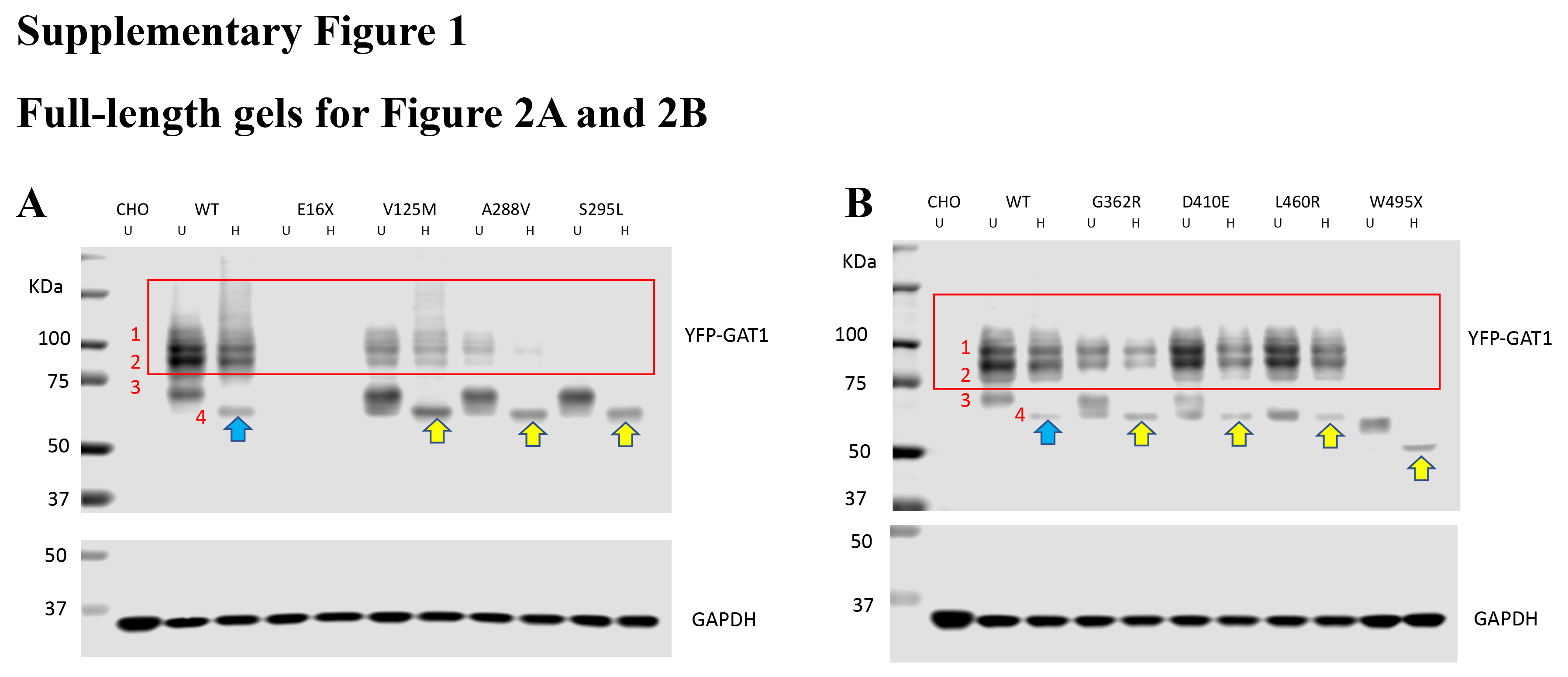

### Supplementary 2

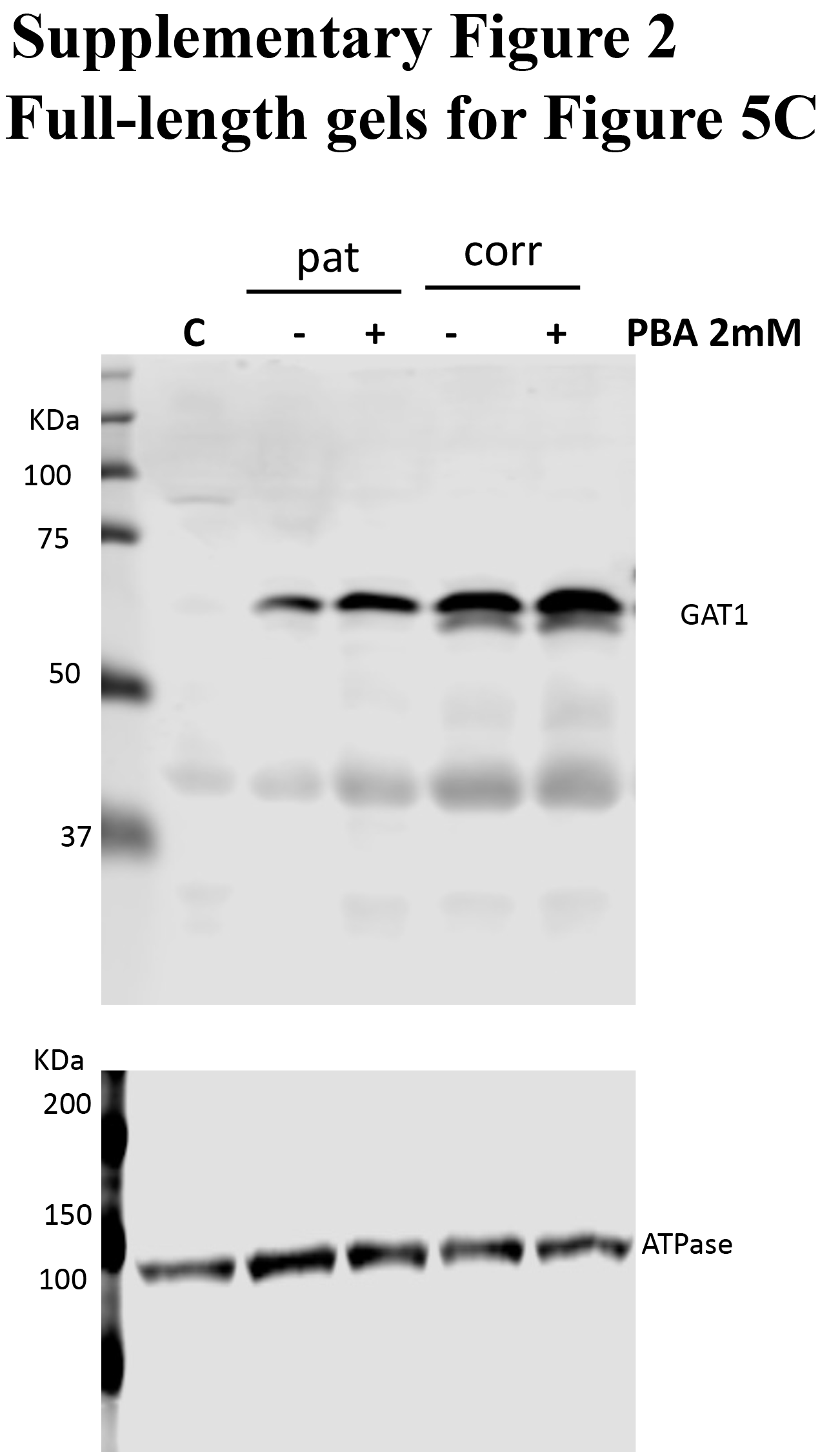

### Supplementary 3

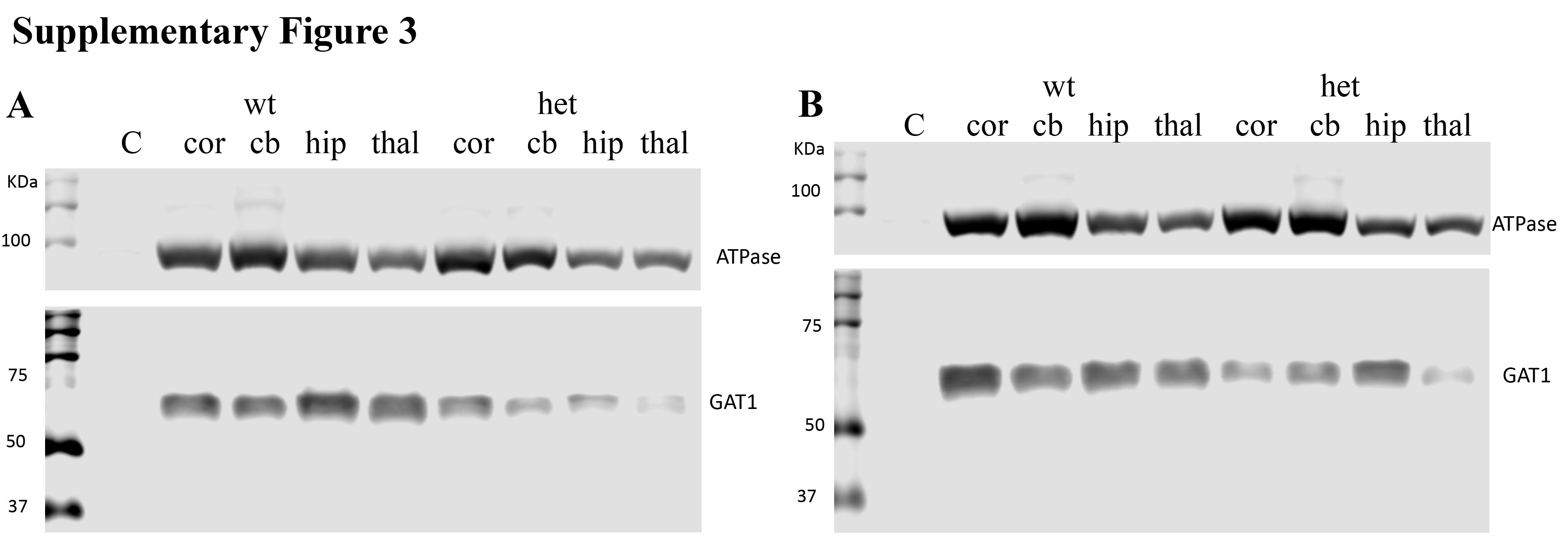

### Supplementary 3B

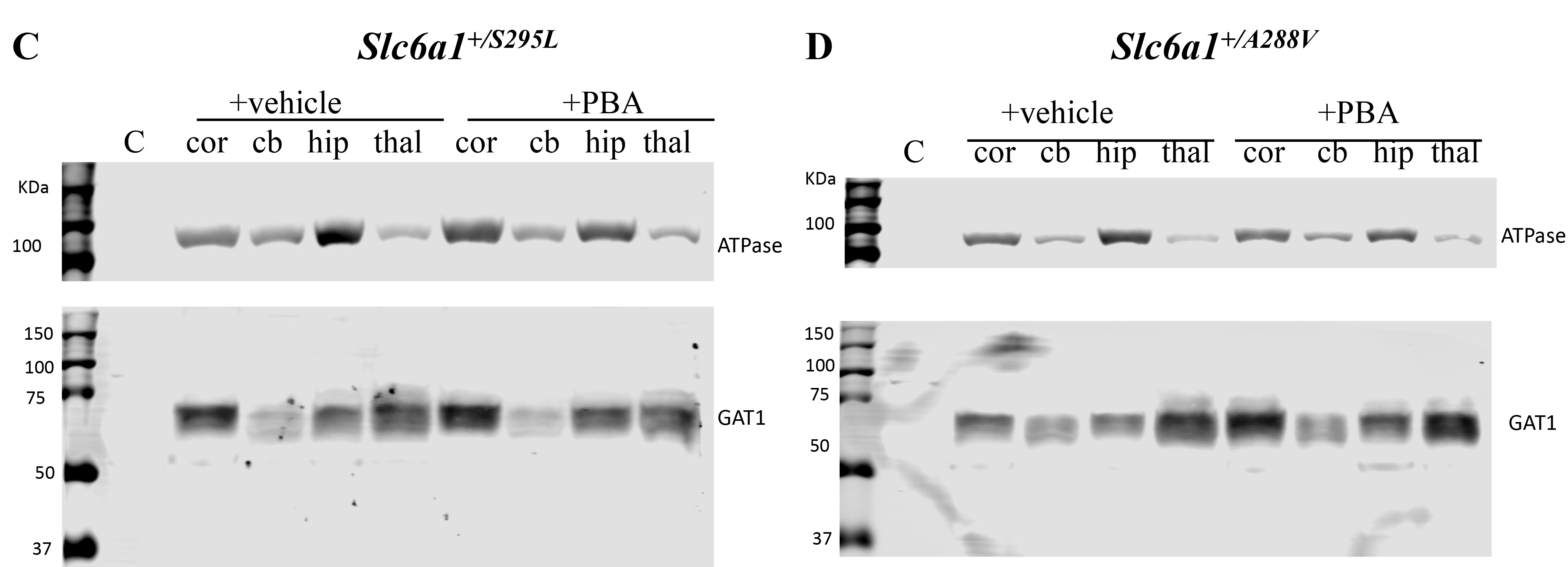

### Supplementary 4

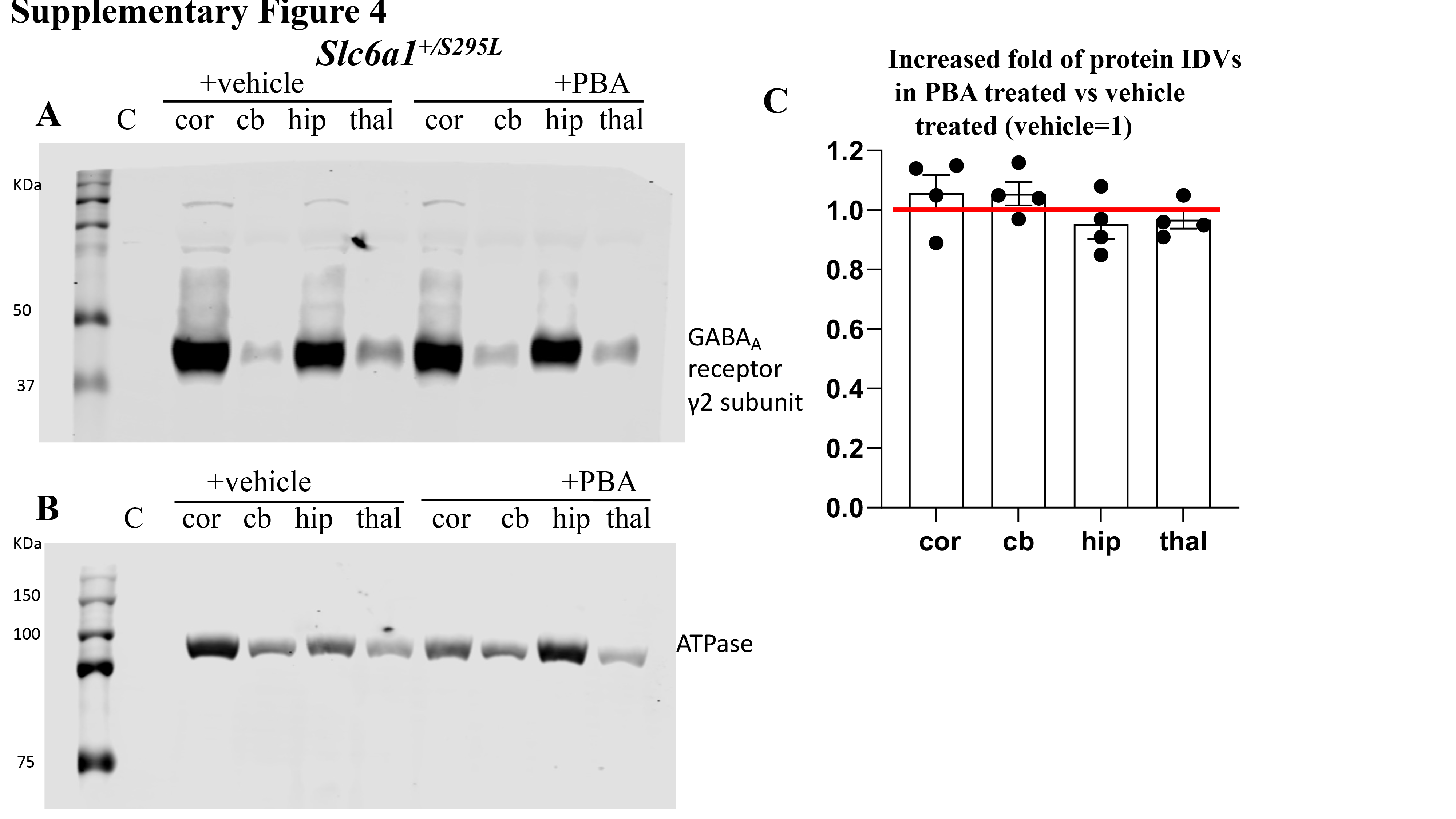
